# Immunogenicity and protective efficacy of an intranasal neuraminidase-based influenza virus vaccine adjuvanted with bacterial cell membrane-derived adjuvants

**DOI:** 10.1101/2025.02.26.640278

**Authors:** Kirill Vasilev, Irene Hoxie, Eduard Puente-Massaguer, Joshua Yueh, Disha Bhavsar, Maya Singh, Corey P. Mallett, Joseph Zimmermann, Florian Krammer

## Abstract

Influenza virus neuraminidase (NA) has emerged as a promising vaccine candidate due to its relatively stable antigenic structure and the ability of NA-specific antibodies to provide cross-protection within influenza virus subtypes. Since the influenza virus causes respiratory infections in humans, developing mucosal vaccines to protect the entry site of the virus is of high importance. Recombinant NA requires adjuvants to induce a protective immune response after mucosal administration. In the current study, we analyze the immunogenicity and protective efficacy of a recombinant NA-based influenza virus vaccine administered intranasally in combination with adjuvants consisting of outer membrane proteins from *Neisseria meningitidis* complexed with exogenous lipopolysaccharides (LPS) from *Shigella flexneri* or endogenous LPS from *N. meningitidis*. We evaluated the local and systemic humoral and cellular immune responses to adjuvanted recombinant N1 NA, analyzing the dynamics of local follicular T-helper (Tfh) cells and germinal center B cells (GCB) in nasal-associated lymphoid tissue (NALT) and tissue-resident memory T cells in lungs, as well as the levels of IgA and IgG in the upper and lower respiratory tracts. Finally, we performed a heterologous challenge study to test the ability of the investigated vaccine formulations to induce cross-protection. The study demonstrates that bacterial cell membrane-derived adjuvants significantly improve the immunogenicity and protective efficacy of the recombinant N1 NA-based influenza vaccine leading to protection against clade 2.3.4.4b H5N1 challenge. This finding supports the potential of these adjuvanted vaccines in providing effective mucosal immunity against influenza virus.

## 1. Introduction

Current influenza virus vaccines elicit narrow, strain-specific immune responses that primarily target the immunodominant head domain of the viral hemagglutinin (HA). While HA immunity may provide superior protection against homologous viral challenge, boosting immunity against the second influenza virus glycoprotein, neuraminidase (NA), is known to be critical for protection and reduction in disease severity when challenged with drifted or heterologous strains. Due to its critical enzymatic functions in the virus life cycle, NA harbors more conserved epitopes^1,2^, thus antibodies against NA are often broadly cross-reactive, and an established correlate of protection, shown to help reduce viral shedding in both infected individuals and animals^3,4^. NA immunity has been associated with reductions in viral transmission in guinea pigs^5,6^ and humans independent of individuals’ pre-existing HA-immunity^7^.

While natural infection induces strong NA-specific immune responses, currently available vaccines are variably effective at inducing robust NA immunity^1^. Vaccines designed to target NA immunity may incidentally help to mitigate the severity of rarer but potentially highly pathogenic, zoonotic subtypes not represented in the current seasonal influenza vaccines, such as avian H5N1, H5N2, H6N1, H7N2, H9N2^8^, as well as non-human mammalian strains within the targeted subtypes, with high zoonotic potential (e.g., swine H3N2, H1N1, canine H3N2). The N1 antigen may be particularly useful for providing protection against avian H5N1^9^, a subtype for which current vaccines inducing HA immunity against H1N1, H3N2 and influenza B virus provide little cross-protection. Cross-reactive, protective epitopes conserved within the N1 subtype have been characterized by monoclonal antibody studies^10–14^ and cross-protection against H5N1 induced by H1N1 exposure has also been well-documented^15–20^. Vaccination studies in humans^21^ and animals have also demonstrated the NA-associated partial cross-protection against seasonal H1N1, 2009 pdmH1N1 and H5N1 viruses^22–24^.

Intranasal vaccination against respiratory pathogens has been demonstrated to provide superior cross-protection compared with intramuscular or intradermal routes of administration^25,26^. Several NA-based influenza virus vaccine studies in animals also demonstrated intranasal immunization can enhance breadth of protection^27–29^, and importantly, provide superior viral transmission inhibition^5,6^. Mucociliary clearance in the airway epithelium is a critical first line of host defense against influenza virus infection. Intranasal vaccination has the potential to induce immune responses from localized systems at the site of initial influenza virus infection involving mucosa-associated lymphoid tissues (MALT), which can help limit infection and transmission. Intramuscular vaccines cannot induce these mucosal immune responses efficiently, so are unable to protect against infection in the upper airway. Intranasal immunization can elicit high antibody titers, in particular mucosal IgA as well as strong cellular immunity, including the promotion of tissue-resident memory T cells. Inducing a strong mucosal immune response against NA would be particularly effective in preventing viral transmission. Inhibition of NA sialidase activity blocks the virus from moving through the mucosa and prevents viral egress, leading to virus aggregates and inhibiting HA-mediated cell entry^30–33^. Notably, intranasal vaccination with recombinant NA (rNA) has also been demonstrated to be more protective against even homologous viral challenge, than recombinant HA (rHA)^34^, supporting the particular value of NA-specific immunity in the mucosa.

Developing vaccination strategies that specifically enhance tissue-resident memory T cells (T_RM_) is important. While both virus-specific antibodies and T_RM_ are necessary for optimal immunity against different strains, T-cell-based protection against IAV relies on the presence of TRMs in the airways and lungs^35–39^. The addition of immunostimulatory adjuvants to recombinant protein vaccination can induce a strong antigen-specific antibody response as well as CD8^+^ and CD4^+^ T-cell responses and elicit T_RM_ cells^40–42^. Adjuvants are also essential for the feasibility of large-scale rNA administration, to lower the effective dose of recombinant protein needed to provide protection^43^. Co-administration of appropriate mucosal adjuvant formulations is needed to effectively penetrate the mucosa and exert a comprehensive series of immune responses^44,45^. As such, the choice and development of safe adjuvants which work effectively in the mucosa, has been a major obstacle for advancing intranasal vaccines^46^.

A 1995 human study demonstrated that NA immunity and vaccine protection could be safely enhanced by supplementing influenza vaccines with soluble NA. However, at that time, NA isolation approaches were not feasible for large-scale production^47,48^. With the recent advancements in rNA protein expression and production, stable, tetrameric rNA with an adjuvant, could now be used as a supraseasonal vaccine to promote NA immunity, offering broad protection across multiple seasons and against drifted strains^49^. This approach has been demonstrated to be safe and effective in many animal studies^6,27,50–54^. However, the majority of these studies only evaluated systemic responses to intramuscular administration of rNA, and even fewer have analyzed cellular immune responses directly. We have previously demonstrated that intramuscular vaccination with recombinant protein constructs composed of the globular head domain of NA, stabilized with a measles virus phosphoprotein tetramerization domain (MPP), can induce strong NA-specific immune responses and confer protection against lethal heterologous virus challenge in mice and hamster models^43,55,56^. This study analyzes the immunogenicity and protective efficacy of the vaccine construct based on the recombinant NA with a measles virus phosphoprotein tetramerization domain (rNA-N1-MPP) influenza virus vaccine adjuvanted with bacterial cell membrane-derived adjuvants BDX100 (also known in the literature under the name Protollin) and BDX301. BDX301 incorporates LPS and the combination of the outer membrane proteins (mostly porin proteins) from *Neisseria meningitidis,* a Gram-negative pathogenic species of bacteria. BDX100 is another adjuvant, comprised of *N. meningitidis* outer membrane proteins and LPS from *Shigella flexneri.* Both are Toll-like receptor (TLR2/TLR4) agonists which have been demonstrated to induce robust mucosal immune responses when administered intranasally^57,58^. Specifically, the BDX100-adjuvanted split H5N1 influenza vaccine stimulated mixed Th1/Th2/Th17 response associated with high IgG2a production and enhanced protection against homologous and heterologous challenge in mice^59^. Another study demonstrated the protective efficacy of a recombinant HA influenza virus vaccine administrated intranasally in combination with BDX100^60^. It was also shown that the BDX100-adjuvanted intranasal split influenza vaccine induces higher HAI and microneutralization (MN) titers compared with AS03-adjuvanted split vaccine administrated intramuscularly^61^. The efficacy of this adjuvant was also demonstrated in a ferret model^62^. BDX100 is safe in humans and was tested as an adjuvant for intranasal trivalent influenza vaccine (TIV) in the human viral influenza challenge model. Intranasal vaccination with Protollin-adjuvanted TIV protected subjects against illness associated with A/Panama/2007/1999 (H3N2) infection^58^.

Fewer data are currently available on the BDX301 adjuvant. In a recent study, however, it was shown that intranasal immunization with a BDX301-adjuvanted Severe acute respiratory syndrome coronavirus 2 (SARS-CoV-2) spike protein induces a superior mucosal immune response compared with alum-adjuvanted formulations^63^. The immunogenicity of NA-based vaccines in combination with adjuvants BDX100 or BDX301 adjuvants was never investigated.

In this study we present a comprehensive analysis of the humoral, cellular, and specifically mucosal immune responses induced by intranasal vaccination with an adjuvanted NA-MPP vaccine. We demonstrate that intranasal administration of rNA-N1-MPP (based on the N1 from A/Michigan/45/2015 (H1N1)) adjuvanted with TLR2/TLR4 agonists, BDX301 or BDX100 promotes robust NA-specific mucosal and systemic immune responses. Mice vaccinated with adjuvanted rNA-N1-MPP showed robust IgA and IgG responses in the upper and lower respiratory tracts and had high NA-inhibiting antibody titers in their sera. Adjuvanted N1-MPP vaccination promoted Tfh and GCB proliferation in NALTs, and CD4^+^ and CD8^+^ T_RM_s in lungs. Notably these robust immune responses were achieved when the vaccine was administered at 5 µl/nare per mouse, thus restricting vaccination to the nasal cavity. Importantly, the adjuvanted rNA-N1-MPP vaccine candidates demonstrated cross-reactive cellular and humoral responses against pre-pandemic seasonal H1N1, 2009 pandemic H1N1, and avian H5N1 viruses. Intranasal administration of adjuvanted rNA-N1-MPP conferred partial protection against the lethal avian H5N1 virus challenge, supporting the potential of targeting NA to enhance the breadth of protection.

## 2. Material and methods

### 2.1. Viruses and cells

A/Michigan/45/2015 (H1N1), A/Singapore/GP1908/2015 (H1N1), A/Netherlands/602/2009 (H1N1)pdm09, A/New Caledonia/20/1999 (H1N1) and A/bald eagle/FL/W22-134-OP/2022 (H5N1 reassortant with A/Puerto Rico/8/1934 vaccine backbone) influenza viruses were used for NA inhibition assay (NAI) and mouse challenge studies. Viruses were grown in 10-day-old embryonated chicken eggs (Charles River Laboratories) at 37 °C for 48 hours with subsequent cooling at 4 °C overnight (O/N). Cell debris was removed by centrifugation of the alantoic fluid at 4000 x *g*, 4 °C, 20 min. Samples were aliquoted and stored at −80 °C. Viral titers were determined by the plaque assay method on Madin-Darby canine kidney (MDCK) cells. MDCK cells were maintained in Dulbecco’s modified Eagle’s medium (DMEM; Gibco) containing 10% fetal bovine serum (FBS; Gibco), 1% penicillin/streptomycin (100 U/ml penicillin, 100 μg/ml streptomycin; Gibco).

### 2.2. Mouse experiments

All experiments with mice were performed following the protocols approved by the Icahn School of Medicine at Mount Sinai Institutional Animal Care and Use Committee (IACUC). Female 6–8-week-old BALB/c or C57/B6 mice (The Jackson Laboratory) were immunized twice with 3 µg of rNA-N1-MPP vaccine alone or in combination with 15 µg of BDX100 or BDX301 adjuvants in a volume of 10 µl (5 µl/nare). Ovalbumin (OVA) was used as an irrelevant control antigen to test the nonspecific protective efficacy of the adjuvants without the vaccine. Challenge infections were performed via intranasal administration of 50 µl of H1N1 A/Singapore/GP1908/2015 (H1N1), A/New Caledonia/20/1999 (H1N1) or A/bald eagle/FL/W22-134-OP/2022 (H5N1) at 5 × 50% lethal dose (LD_50_) for survival analysis and body weight monitoring or 0.1 × LD_50_ for viral titers analysis. For the intranasal administration (vaccinations and challenges), mice were anesthetized with 100 µl of ketamine-xylazine cocktail (87.5 mg/kg, 12.5 mg/kg) administrated intraperitoneally. The volumes of the vaccines and viruses administrated intranasally were distributed equally between the nares in a drop-by-drop manner. Mice were unconscious during the procedures. Euthanasia was performed by cervical dislocation following the approved IACUC protocol. Experiment designs are outlined in Fig. 1a-d.

**Figure 1.**
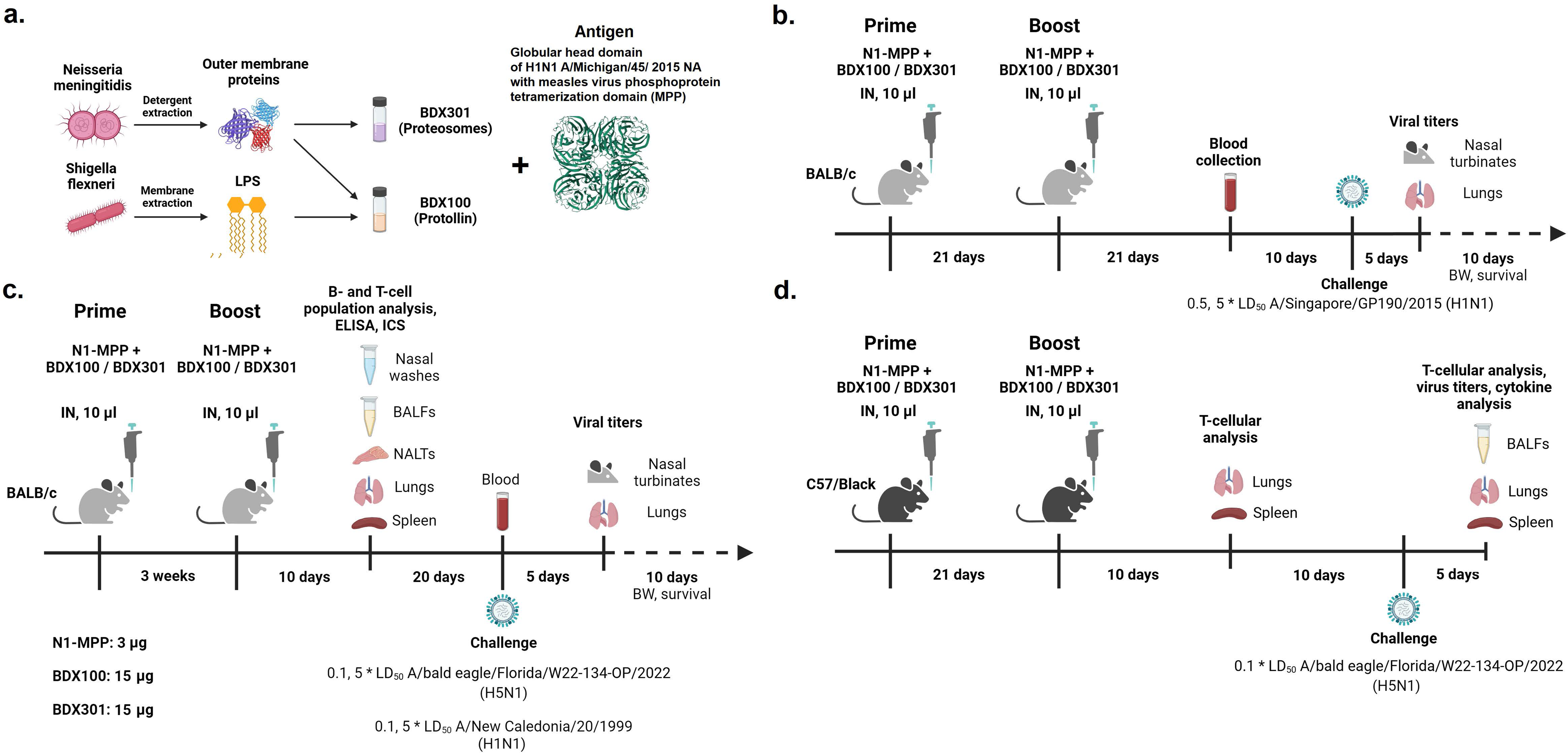
Outline of the study design. **a.** Adjuvants and vaccine preparation outline **b.** Homologous protection study scheme. BALB/c mice were intranasally vaccinated twice with rNA-N1-MPP adjuvanted with BDX100 or BDX301 adjuvants in a volume of 10 µl (5 µl/nare). Ovalbumin (OVA) was used as an irrelevant control antigen. Four weeks after the second immunization, mice were challenged with 5 × LD_50_ of homologous influenza strain A/Singapore/GP1908/2015 H1N1. The body weight and survival of the experimental animals were monitored for two weeks after the challenge. **c.** Immunogenicity and heterologous protection efficacy study outline. On day 10 after the boost nasal washes, BALFs, NALTs, lungs, and spleens were collected for the immunological read-outs. Heterologous influenza virus strains (A/bald eagle/Florida/ W22-134-OP/2022 (H5N1 reassortant with A/Puerto Rico/8/1934 vaccine backbone) and A/New Caledonia/20/1999 (H1N1)) were used for the challenge study. **d.** T-cellular immune response analysis study outline. C57/B6 mice were used to assess the T-cellular immune response in lungs and spleens to the rNA-N1-MPP vaccine adjuvanted with BDX100 or BDX301 10 days after the booster immunization and 5 days after the heterologous challenge with A/bald eagle/Florida/W22-134-OP/2022 (H5N1) influenza virus strain.

### 2.3. Plaque assay

To assess viral titers samples were serially diluted 1:10 in minimal essential medium (MEM). Diluted samples were incubated for 1 hour with MDCK cells seeded at a density of 3 × 10^5^ cells per well on the previous day. An agarose overlay was applied to the cells at the end of the incubation period. The overlay included 0.64% (w/v) agarose (Oxoid) in MEM supplemented with 2 mM l-glutamine, 0.1% (w/v) sodium bicarbonate, 10 mM 2-(4-(2-hydroxyethyl)piperazin-1-yl)ethanesulfonic acid (HEPES), penicillin (100 U/ml), streptomycin (100 μg/ml), 0.2% (w/v) bovine serum albumin (BSA), L-(tosylamido-2-phenyl) ethyl chloromethyl ketone (TPCK)- treated trypsin (1 μg/ml), and 0.1% (w/v) diethyl aminoethyl-dextran. Plates were incubated at 37 °C for 72 hours. Plaques were counted after fixation with 3.7% (v/v) formaldehyde in PBS, followed by visualization through immunostaining. Mouse antisera and sheep anti-mouse IgG (H&L) peroxidase conjugate (Rockland) were used for the plaque staining. All virus titers are reported in units of log_10_ plaque-forming units (PFU) per milliliter.

### 2.4. Recombinant proteins and adjuvants

Recombinant N1 constructs utilized in these experiments were produced in insect cells using the baculovirus expression system, as described in prior studies^55,64^. Briefly, the rNA-N1-MPP construct’s coding sequence was inserted into a modified pFastBac vector. This vector included an N-terminal signal peptide, a hexahistidine purification tag (His-tag), a measles virus phosphoprotein (MPP) tetramerization domain, a thrombin cleavage site, and the globular head domain of N1 derived from the A/Michigan/45/2015 (H1N1) influenza virus strain. The recombinant pFastBac constructs were introduced into DH10Bac bacteria (Invitrogen). Transformed colonies were selected, screened, and cultured for the extraction of bacmid DNA using the PureLink Plasmid Filter Midiprep Kit (Invitrogen). This bacmid DNA was then transfected into Sf9 cells with Cellfectin II (Invitrogen) to rescue recombinant baculovirus.

Western blot analysis was used to verify the protein identity. The rescued baculovirus was amplified in Sf9 cells and subsequently used to infect High Five cells for protein production. Proteins were harvested from the culture supernatant and purified through Ni²⁺–nitrilotriacetic acid resin chromatography as previously described^65^. Protein concentration was determined using Quick Start Bradford 1× Dye Reagent (BioRad), and purified proteins were stored at −80 °C in PBS.

Sodium dodecyl-sulfate polyacrylamide gel electrophoresis (SDS-PAGE) was performed using 4–20% polyacrylamide mini PROTEAN TGX gels (Bio-Rad) to assess protein integrity and tetramerization. N1 proteins (A/Puerto Rico/8/1934 N1, A/New Caledonia/20/1999 N1, A/Wisconsin/67/2022 N1, A/Vietnam/1203/2004 N1, A/bald eagle/FL/W22-134-OP/2022 N1, A/dairy cattle/Texas/24-008749-001-original/2024 N1) used for enzyme-linked immunosorbent assays (ELISAs), were produced using a similar protocol as the N1-MPP protein but construct contained a vasodilator-stimulated phosphoprotein (VASP) tetramerization domain instead of the MPP domain to minimize cross-reactivity due to antibodies targeting the tetramerization domain. BDX100 and BDX301 adjuvants were prepared as described previously^66^.

### 2.5. ELISA

For the ELISA, 96-well Immulon 4HBX plates (Thermo Fisher Scientific) were coated with recombinant proteins at a concentration of 2 μg/ml in PBS (pH 7.4), with 50 μl added per well. The plates were incubated overnight at 4 °C. The next day, the plates were washed three times using PBS containing 0.1% (v/v) Tween 20 (PBS-T). Blocking was performed with a 3% (v/v) goat serum and 0.5% (w/v) nonfat dry milk solution in PBS-T for one hour at room temperature. After blocking, mouse serum was added to the first well at a 1:30 dilution (150 μl per well) and serially diluted 1:3. The plates were incubated for two hours at 20 °C, followed by another three washes with PBS-T. To detect total IgG, a 1:3000 dilution of sheep anti-mouse IgG (H&L) conjugated to peroxidase (Rockland) in blocking solution was applied. For color development, 100 μl of O-phenylenediamine dihydrochloride (OPD) substrate (SigmaFast OPD, Millipore Sigma) was added to each well. After a 10-min incubation, the reaction was terminated by adding 50 μl of 3 M hydrochloric acid (HCl) per well. Optical density at 490 nm (OD490) was measured using a Synergy H1 microplate reader (BioTek).

### 2.6. Neuraminidase inhibition assay (NAI)

The NAI assay was performed to evaluate NA-inhibiting antibody titers in serum samples against A/Michigan/45/2015 (H1N1), A/Singapore/GP1908/2015 (H1N1), and A/Netherlands/602/2009 (H1N1)pdm09 influenza viruses, following an established protocol^67^. Briefly, 96-well Immulon 4HBX plates (Thermo Fisher) were coated with 100 μl/well of fetuin solution (50 μg/ml; Sigma Aldrich) and incubated overnight at 4 °C. Serum samples were heat-inactivated at 56 °C for one hour and 10 serial twofold dilutions were prepared across the 96-well plate starting from 1:15 dilution. Wells #11 were left as virus-only control (no serum was added) and wells #12 were left as blanks. The diluted serum samples were mixed with an equal volume of twice the 50% effective concentration (EC_50_) of each virus and incubated for two hours at room temperature with shaking. The fetuin-coated plates were washed three times with PBS containing 0.1% Tween 20 (PBS-T), and 100 μl of the serum-virus mixtures were transferred to the fetuin-coated wells. The plates were then incubated overnight at 37 °C. After incubation, the plates were washed three times with PBS-T and treated with 100 μl/well of horseradish peroxidase (HRP)-conjugated peanut agglutinin (PNA). The plates were incubated for one hour at room temperature and washed six times. Color development reaction and data collection were performed as described for the ELISA. The percentage of NA inhibition for each well was calculated using the formula: %inhibition = (1 - OD490_sample_ / OD490_virus only_) × 100. The 50% inhibitory dilution (ID_50_) was calculated in RStudio (R version 4.3.3, RStudio Inc).

### 2.7. Flow cytometry

#### 2.7.1. Tissue preparation

Mice were humanely euthanized, and their chest cavities were opened. To collect the broncho-alveolar lavage fluids (BALFs) the lungs were instilled with 1 ml of cold PBS, and the fluid was aspirated. Subsequently, the right ventricle was perfused with 10 mL of cold PBS. Lungs were harvested and processed using a gentleMACS Octo Dissociator for tissue homogenization and a Lung Dissociation Kit (Miltenyi Biotec) to obtain a single-cell suspension. The cell suspension was filtered through a 70-μm cell strainer (BD Biosciences) after the digestion. Spleens were mechanically disrupted using pestle homogenizers and filtered similarly. NALT samples were collected as described previously^68^. Lymphoid tissue was scraped from the upper palate and mechanically disrupted. Erythrocytes were lysed using red blood cell (RBC) lysis buffer (BioLegend) and the reaction was stopped with 2% fetal bovine serum (FBS) in PBS. The resulting cell suspension was centrifuged, and the pellet was resuspended in complete Roswell Park Memorial Institute (RPMI) 1640 media. For T-cell immune response analysis, 2 × 10^6^ cells from lung and spleen samples were seeded in 96-well plates for antigen-specific stimulation. Immune cell populations in the lung, spleen, and NALT samples were stained with fluorescently labeled antibodies right after the tissue processing. BALFs were centrifuged at 4000 rpm for 5 min to remove cells and debris, aliquoted, and stored at –80 °C for the viral titer and cytokine analysis.

#### 2.7.2. T-cellular immune response analysis (intracellular cytokine staining (ICS))

To assess antigen-specific T-cell responses, cells were stimulated with a peptide pool spanning the entire sequence of the influenza A/California/04/2009 (H1N1) NA virus protein. The peptide pool consisted of 15-mer peptides with an 11 amino acid overlap, each at a concentration of 5 µg/ml. Cells were stimulated in the presence of Brefeldin A, Monensin, and the costimulatory anti-CD28 antibodies to mimic the secondary signal during the antigen presentation. Negative control samples were incubated without specific antigens. Following stimulation, cells were stained with fluorescently labeled antibodies (0.5 μl per sample of CD3-BV711 (Biolegend), 0.125 μl per sample of CD4-PerCP/Cy5.5 (Tonbo Biosciences), 0.25 μl per sample of CD8-BV785 (Biolegend), 0.125 μl per sample of CD62L-APC/Cy7 (Biolegend), and 0.25 μl per sample of CD44-PE/Cy7 (Tonbo Biosciences)). Intracellular staining of cytokines was performed using the Cytofix/Cytoperm Fixation/Permeabilization Solution Kit (BD Biosciences) following the manufacturer’s instructions. Antibodies for TNFα-AF488 (0.25 μl per sample), IFNγ-BV421 (0.5 μl per sample), and IL-2-PE (0.5 μl per sample) (BioLegend) were used to identify the corresponding cytokines. ZombieAqua (BioLegend) viability dye was used to discriminate between live and dead cells. To perform the group comparison the percentage of cytokine-producing cells in non-stimulated samples were subtracted from the corresponding values for peptide-stimulated samples.

#### 2.7.3. Cell population analysis

For the analysis of T- and B-cell populations in lungs, spleen, and NALTs, cells were stained with the antibody cocktail, including the following antibodies: CD38-AF488 (0.125 μl, BioLegend), PD-1-PE/Dazzle (0.5 μl, BioLegend), CD273-RB744 (0.25 μl, BD Biosciences), CD138-PE (0.25 μl, BioLegend), CD103-PE-Cy5.5 (0.5 μl, BioLegend), CD44-PE/Cy7 (0.25 μl, Tonbo Biosciences), GL7-APC (0.25 μl, BioLegend), CD69-AF700 (1 μl, BioLegend), CD62L-APC/Cy7 (0.125 μl, BioLegend), CD185-BV421 (0.5 μl, BioLegend), CD80-BV605 (1 μl, BioLegend), IgD-BV650 (0.25 μl, BioLegend), CD3-BV711 (0.5 μl, BioLegend), CD8-BV785 (0.25 μl, BioLegend), CD4-BUV496 (0.125 μl, BD Biosciences), IgM-BUV737 (1 μl, BD Biosciences), CD19-BUV805 (0.125 μl, BD Biosciences). Data were collected using FACSymphony A5 SE High-Parameter Cell Analyzer (BD Biosciences). For the analysis of the innate immune cell populations in the lungs, cells were stained with the following antibody cocktail: CD117-AF488 (0.25 μl, BioLegend), Ly6C-PerCP/Cy5.5, (0.25 μl, BioLegend), CD103-APC (1 μl, BioLegend), Ly6G-AF700 (0.25 μl, BioLegend), CD11b-APC/Cy7 (0.25 μl, BioLegend), CD64-BV421 (1.5 μl, BioLegend), CD11c-BV605 (1 μl, BioLegend), MHC-II (IA/IE)-BV650 (0.25 μl, BioLegend), CD24-BV711 (0.25 μl, BioLegend), CD45-BV785 (0.125 μl, BioLegend), FCeRI-PE (0.5 μl, BioLegend) SiglecF-PE/CF594, (0.5 μl, BD Biosciences). Data were collected using an Attune flow cytometer (Thermo Fisher Scientific). FlowJo_v10.10.0 and RStudio (R version 4.3.3.) software were used for the flow cytometry data analysis and visualization.

#### 2.7.4. Cytokine analysis

A LEGENDplex kit (BioLegend) was used to quantify proinflammatory and Th1/Th2/Th17-associated cytokines in BALFs. The assay was performed following the manufacturer’s instructions. Briefly, samples were incubated with capture beads specific to the target cytokines. After washing, biotinylated detection antibodies were added following the addition of streptavidin-conjugated phycoerythrin (PE). Data were collected using an Attune flow cytometer (Thermo Fisher Scientific) and analyzed with LEGENDplex data analysis software (https://legendplex.qognit.com).

### 2.8. Statistical analysis

Group comparisons were performed using one-way analysis of variance (ANOVA) with Tukey’s correction for multiple comparisons in RStudio software (R version 4.3.3, RStudio Inc). Comparison with a single control group was performed using Dunnett’s test. Differences with P-values equal or less than 0.05 were considered statistically significant. Uniform Manifold Approximation and Projection (UMAP) v4.1.1 and PhenoGraph v4.0.5 plugins for FlowJo were used for dimensionality reduction and clusterization of the flow cytometry data. UMAP results were visualized using R. Fluorescence data were transformed using square root transformation and normalized using min-max normalization.

## 3. Results

### 3.1. rNA-N1-MPP vaccine adjuvanted with BDX100 or BDX301 adjuvants protects mice from the challenge infection with A/Singapore/GP1908/2015 H1N1 influenza virus

To evaluate the protective efficacy of the rNA-N1-MPP influenza vaccine formulated with BDX100 or BDX301 adjuvants, mice were intranasally immunized twice at 21 days interval with 3 µg of rNA-N1-MPP combined with 15 µg of either BDX100 or BDX301 in a 10 µl volume (5 µl per nare). Ovalbumin (OVA) was used as an irrelevant control antigen (Fig. 1a–c). On day 10 after the second immunization, lungs, spleens, and nasal-associated lymphoid tissues (NALTs) were collected to assess immune cell population dynamics. Both BDX100 and BDX301 induced a notable increase in the percentage of Tfh cells in the spleens. Additionally, a trend toward higher levels of activated and memory B cells was observed in the adjuvanted groups, particularly in mice receiving the BDX301-adjuvanted vaccine (Fig. 2a).

**Figure 2.**
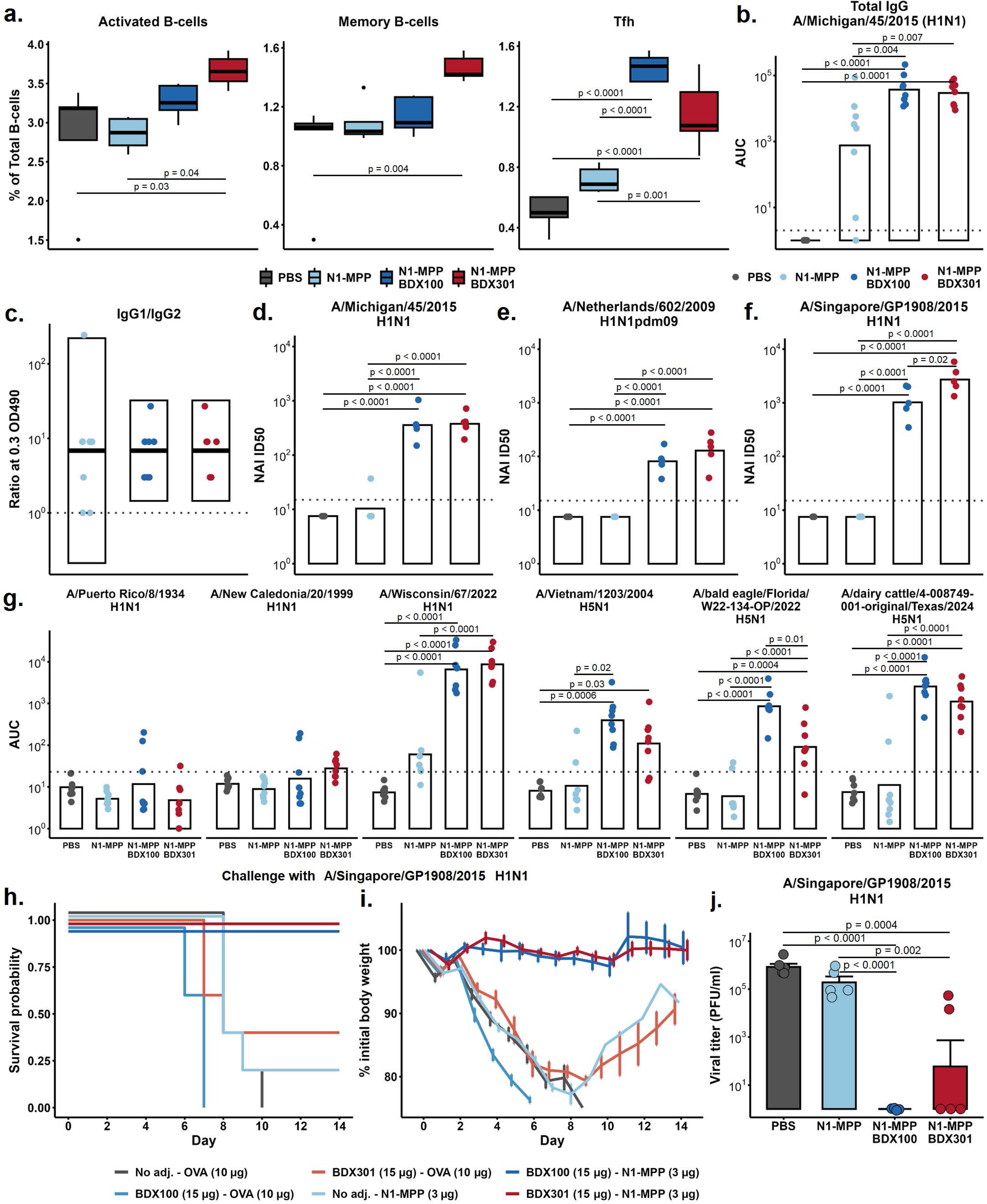
Systemic immune response and protection efficacy of rNA-N1-MPP vaccine adjuvanted with BDX100 or BDX301 adjuvants. **a.** B-and follicular T-helper (Tfh) cell subpopulation percentages (activated B-cells: CD3^‒^CD19^+^CD38^+^GL7^‒^CD273^‒^CD80^+^, memory B-cells: CD3^‒^CD19^+^CD38^+^GL7^‒^CD273^+^CD80^+^, Tfh: CD3^+^CD4^+^CD185^+^PD1^+^) in the spleens of vaccinated mice 10 days post boost (n = 5). **b.,e.,f.** Neuraminidase inhibition activity of sera from the vaccinated mice against various influenza virus strains (A/Michigan/45/2015 H1N1, A/Singapore/GP1908/2015 H1N1, A/Netherlands/602/2009 (H1N1)pdm09). Dots represent individual ID_50_ values for each sample (n = 5). Bars represent average values. **c.** Total IgG response against the recombinant neuraminidase protein from homologous influenza virus strain A/Michigan/45/2015 H1N1 measured with ELISA. Dots represent individual areas under curves (AUC) values (n = 8). **d.** IgG1/IgG2a ratio at 0.3 OD490. Dots represent individual values (n = 8). **g.** Total IgG response against the recombinant neuraminidase proteins from various heterologous influenza virus strains (A/Puerto Rico/8/1934 H1N1, A/New Caledonia/20/1999 H1N1, A/Wisconsin/67/2022 H1N1, A/Vietnam/1203/2004 H5N1, A/bald eagle/FL/W22-134-OP/2022 H5N1, A/dairy cattle/Texas/24-008749-001-original/2024 H5N1). Dots represent individual AUC values; bars represent average values (n = 8). **h. i.** Survival and body weight dynamics after the challenge with 5 × LD_50_ of A/Singapore/GP1908/2015 H1N1 (n = 10). **j.** Viral titers in lungs on day 5 after the challenge with A/Singapore/GP1908/2015 H1N1 (n = 5) **a.-g.,j.** Groups were compared using ANOVA followed by the Tukey posthoc test. P-values for the groups with statistically significant differences are shown on the plots.

The humoral immune response was assessed on day 21 post-immunization via NAI and ELISA against homologous and heterologous influenza virus strains (NAI) and NA-proteins (ELISA). Both BDX100 and BDX301 significantly increased total anti-NA IgG levels compared to the nonadjuvanted rNA-N1-MPP group (Fig. 2b). The IgG1/IgG2a ratio remained skewed toward IgG1 across all groups, indicating a predominant Th2 response (Fig. 2c). The adjuvants enhanced the production of anti-NA antibodies with NAI activity compared to the non-adjuvanted vaccine (Fig. 2d–f). To assess cross-reactivity, sera were tested against a panel of NA antigens from various influenza virus strains. No or little cross-reactivity was observed with A/Puerto Rico/8/1934 H1N1 or A/New Caledonia/20/1999 H1N1 NAs. However, sera from mice immunized with adjuvanted rNA-N1-MPP exhibited cross-reactivity with more recent variants, including A/Wisconsin/67/2022 H1N1, A/Vietnam/1203/2004 H5N1, A/bald eagle/FL/W22-134-OP/2022 H5N1, and A/dairy cattle/Texas/24-008749-001-original/2024 H5N1 NA. The non-adjuvanted vaccine did not demonstrate cross-reactivity against these strains (Fig. 2g). Notably, rNA-N1-MPP with BDX100 induced a higher immune response to A/bald eagle/FL/W22-134-OP/2022 H5N1 NA compared to BDX301, though antibody responses to other antigens were similar between the two adjuvanted groups.

Three weeks after the second immunization, blood samples were collected via cheek bleeding for humoral immune response analysis. Four weeks post-immunization, mice were challenged with 5 or 0.5 × LD_50_ of the homologous H1N1 influenza virus strain A/Singapore/GP1908/2015. Body weight and survival were monitored for two weeks post-challenge with the 5 LD_50_ dose (Fig. 2h–i). On day 5, lungs were collected from mice challenged with 0.5 × LD_50_ to evaluate viral replication (Fig. 2k). Mice receiving the non-adjuvanted vaccine, adjuvants with an irrelevant antigen (OVA), or PBS showed rapid weight loss after the challenge. In the PBS group, all animals reached the humane endpoint and were sacrificed by day 10. The nonadjuvanted rNA-N1-MPP vaccine failed to protect mice from challenge infection. While BDX100 alone did not improve survival or body weight compared with controls, BDX301 alone slightly increased survival, though without statistically significant effects on weight loss. In contrast, mice vaccinated with rNA-N1-MPP adjuvanted with either BDX100 or BDX301 were fully protected from mortality and maintained stable body weight post-challenge. rNA-N1-MPP with BDX100 induced complete protection from the A/Singapore/GP1908/2015 influenza virus replication in the lungs on day 5 after the challenge. In the BDX301-adjuvanted group, most vaccinated animals demonstrated full protection from viral replication. In two animals the vaccination induced infection-permissive immunity, but the viral titers were lower compared to non-adjuvanted and PBS controls (Fig. 2j).

### 3.2. BDX100 and BDX301 adjuvants enhance the mucosal immune response to the rNA-N1-MPP vaccine in the upper respiratory tract

To evaluate the effect of the adjuvants on the local mucosal immune response in the upper respiratory tract (URT), nasal washes and NALTs were collected on day 10 following the second booster vaccination (Fig. 1c). The analysis of immune cell dynamics in the NALTs utilized an 18-color fluorescent antibody panel targeting key markers of mouse T and B lymphocytes. The most notable changes in immune cell subpopulations were observed for the GCB and Tfh (Fig. 3a, b). Both BDX100 and BDX301-adjuvanted vaccine formulations significantly increased the percentages of Tfh and GCB cells compared with the non-adjuvanted vaccine. These increases were associated with enhanced production of IgA and IgG in the nasal washes, although no statistically significant correlation was found between the cellular changes and antibody levels (Fig. 3c). For a detailed characterization of lymphocyte subpopulations in the NALTs, the PhenoGraph clustering algorithm was applied to pooled sample data. Differential expression of phenotypic markers and average cluster percentages are illustrated in Fig. 3e, f, i. A total of 20 clusters were identified based on the 18 analyzed markers. Statistical analyses of cluster differences compared with the PBS control group are presented in Fig. 3h. The most significant differences between vaccinated and non-vaccinated (PBS) groups were shown for Cluster 2, characterized by Tfh markers CD185 and PD1, Cluster 12, expressing the GCB marker GL7, Cluster 1, representing CD69^+^CD44^+^ activated T cells. These clusters were elevated in adjuvanted vaccine groups. Additionally, Clusters 18 and 20, defined as IgM^lo^IgD^lo^ and IgM^+^IgD^+^CD69^+^ B cells, respectively, showed increased percentages, suggesting a higher prevalence of isotype-switched and activated B lymphocytes in the adjuvanted groups. Interestingly, Cluster 6, defined by CD4^+^CD44^‒^CD62L^‒^ effector T cells, was elevated exclusively in the group receiving the nonadjuvanted N1-MPP. In contrast, the percentages of these cells in the adjuvanted groups were similar to those in the PBS control. Memory B-cell populations also showed notable changes: Cluster 15, representing CD62L^+^ memory B cells, and Cluster 10, representing CD80^+^CD273^+^ memory B cells, had reduced percentages in vaccinated groups compared with the PBS control (Fig. 3j). This reduction may reflect the reactivation of memory B cells after booster immunization, leading to phenotypic changes or redistribution to other sites.

**Figure 3.**
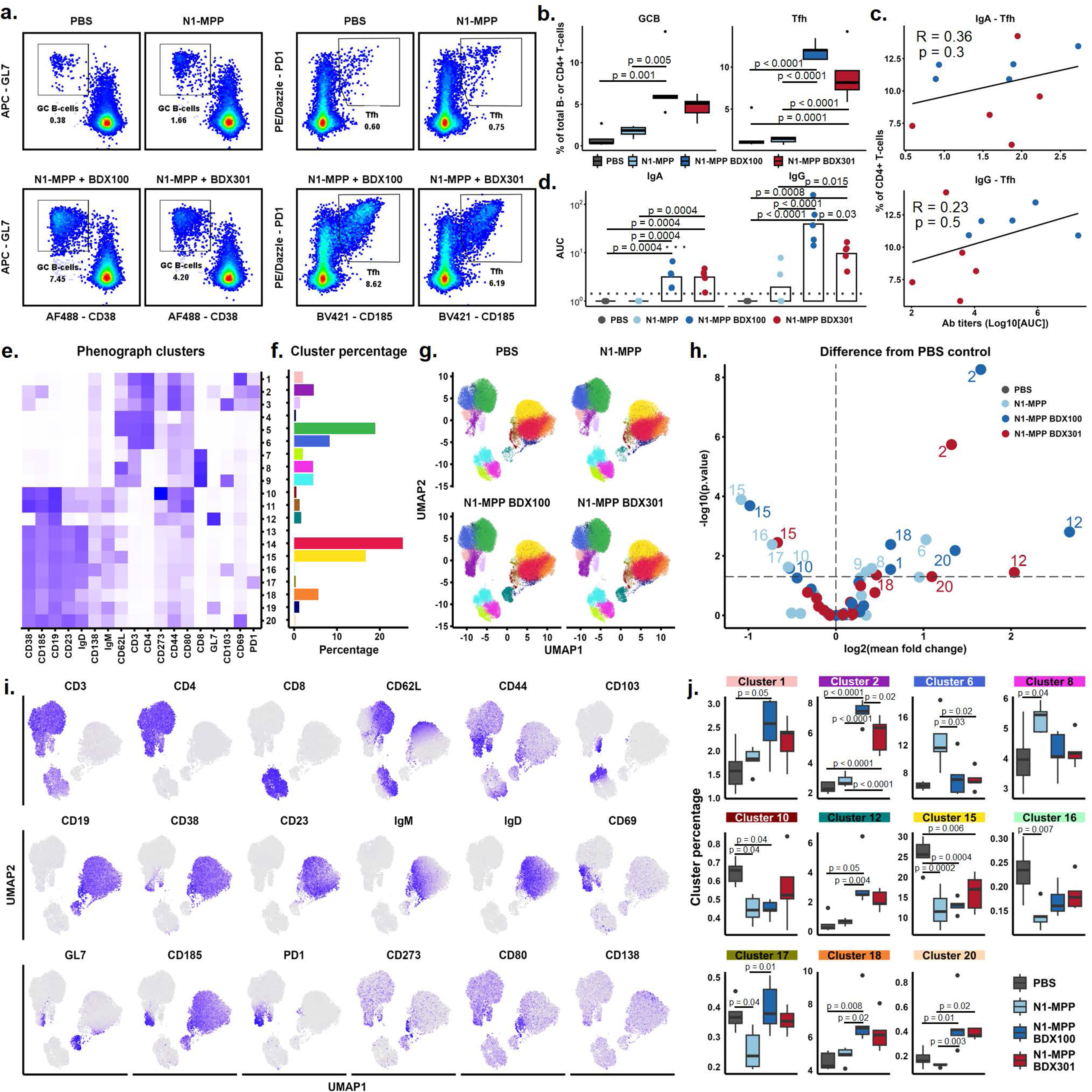
Local immune response in nasal washes and NALTs in mice immunized with rNA-N1-MPP vaccine adjuvanted with BDX100 or BDX301 adjuvants. **a.** Representative flow cytometry plots showing the percentage of GCB add Tfh populations in NALTs (gated on live CD3^‒^CD19^+^ and CD3^+^CD19^‒^CD4^+^ cells, respectively). **b.** Percentage of GCB add Tfh populations in NALTs 10 days after the booster immunization (n = 5). **c.** Pearson’s correlation between the percentage of Tfh in NALTs and the mucosal IgA/IgG levels in nasal washes. **d.** Level of IgA and IgG in the nasal washes obtained from the immunized animals 10 days after the booster immunization. Dots represent individual AUC values; bars represent average values (n = 5). Groups were compared using ANOVA followed by the Tukey posthoc test. P-values for the groups with statistically significant differences are shown on the plots. **e.** Differential expression of phenotype markers in NALT subpopulations identified with PhenoGraph clustering algorithm. **f.** PhenoGraph cluster percentage from the total number of live cells in NALTs 10 days after the booster immunization. Samples were combined for percentage calculation. **g.** UMAP projections showing the distribution of PhenoGraph clusters in NALTs of mice 10 days after the booster immunization with adjuvanted or nonadjuvanted N1-MPP or PBS. **h**. Volcano plot representing the magnitude and significance of PhenoGraph cluster percentage changes compared to the corresponding values in the PBS-control group. **i.** UMAP projections showing the differential expression of phenotype markers on NALT cells. **j.** Percentage of PhenoGraph clusters for which the statistical differences were demonstrated between the experimental groups in the ANOVA test (n = 5). Groups were compared using ANOVA followed by the Tukey posthoc test. P-values for the groups with statistically significant differences are shown on the plots.

### 3.3. BDX100 and BDX301 adjuvants enhance the mucosal immune response to the rNA-N1-MPP vaccine in the lungs

Cell population dynamics and local IgA and IgG antibody responses were assessed in mouse lungs and bronchoalveolar lavage fluids (BALFs) on Day 10 following the booster vaccination. Mice receiving the vaccine supplemented with BDX100 or BDX301 exhibited increased percentages of memory B cells (CD19^+^CD38^+^GL7^‒^CD273^+^CD80^+^) and higher levels of NA-specific IgA and IgG antibodies in BALFs (Fig. 4b–d). Notably, BDX100 induced higher levels of IgA than BDX301, while IgG responses were similar between the two groups. In contrast, the non-adjuvanted rNA-N1-MPP vaccine failed to induce IgA production in the lungs and elicited only a marginal IgG response. BDX100 and BDX301 also enhanced the accumulation of CD4^+^ and CD8^+^ memory T cells in the lungs of immunized mice (Fig. 4e–m). The most significant changes, compared to the rNA-N1-MPP and PBS control groups, were observed in CD4^+^ T lymphocyte subpopulations. The adjuvanted groups, particularly those receiving BDX100, showed elevated percentages of effector-memory (EM; CD4^+^CD44^+^CD62L^‒^) T cells, as well as activated (CD4^+^CD69^+^CD103^‒^) and tissue-resident (CD4^+^CD69^+^CD103^+^) memory T cells. No such changes were observed in the non-adjuvanted group. Although CD8^+^ T-cell subpopulations were also affected by the adjuvants, the changes were less pronounced than those on CD4^+^ T cells. The total percentage of EM CD8^+^ T cells did not significantly differ between the adjuvanted and control groups. However, the dynamics of activated and tissue-resident CD8^+^ memory T cells closely paralleled those of the corresponding CD4^+^ subpopulations (Fig. 4k–m).

**Figure 4.**
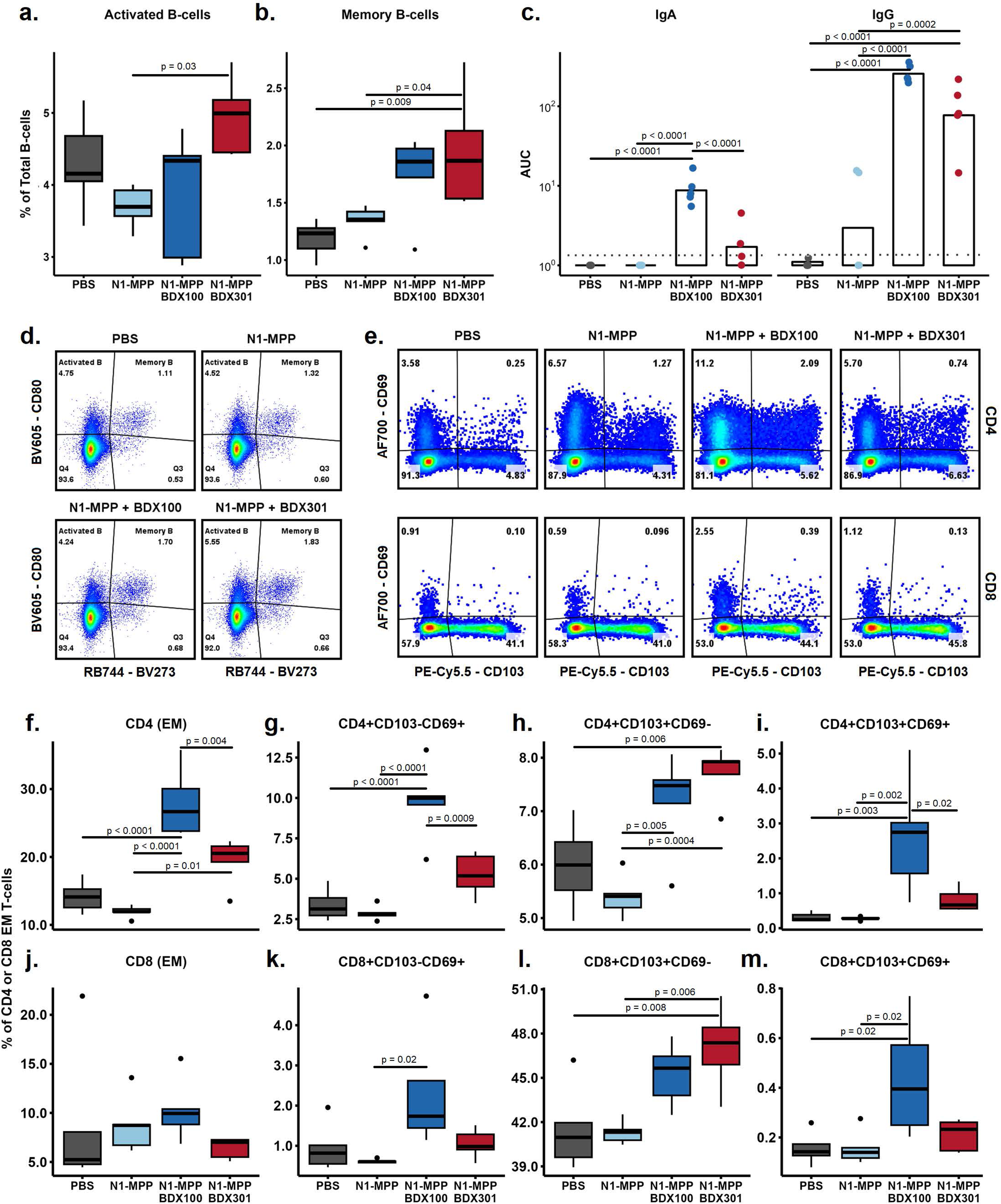
Local immune response in BALFs and lungs in mice immunized with rNA-N1-MPP vaccine adjuvanted with BDX100 or BDX301 adjuvants. **a.** Percentage of activated B-cells and **b.** memory B-cells in the lungs of vaccinated mice 10 days after the booster immunization (n = 5). c. Level of IgA and IgG in the BALFs obtained from the immunized animals. Dots represent individual AUC values (n = 5); bars represent average values. **d.** Representative flow cytometry plots showing the percentage of different B-cell subpopulations in the lungs of immunized mice (gated on live CD3^‒^CD19^+^CD38^+^GL7^‒^ cells). **e.** Representative flow cytometry plots showing the percentage of tissue-resident memory T-cell subpopulations in the lungs of immunized mice (gated on live CD3^+^CD19^‒^CD4/CD8^+^ cells). **f.-m.** T-cell subpopulation percentage in the lungs of mice 10 days after the booster immunization (n = 5). Groups were compared using ANOVA followed by the Tukey posthoc test. P-values for the groups with statistically significant differences are shown on the plots.

To investigate the antigen-specific T-cell immune response, C57BL/6 mice were immunized twice with the rNA-N1-MPP vaccine alone or combined with BDX100 or BDX301 adjuvants (Fig. 1d). Lungs and spleens were collected on Day 10 after the second immunization, and cells were restimulated with overlapping peptides spanning the full NA protein sequence. The strongest T-cell immune response was observed in the group receiving the BDX301-adjuvanted vaccine. This antigen-specific response was primarily mediated by CD4^+^ T lymphocytes, while CD8^+^ T cells did not exhibit a significant increase in cytokine production upon restimulation (SFig. 1). The T-cell immune response was observed in both lungs and spleens, predominantly involving polyfunctional effector-memory (IFNγ^‒^IL2^+^TNFα^+^) CD4^+^ T cells (Fig. 5a, b).

**Figure 5.**
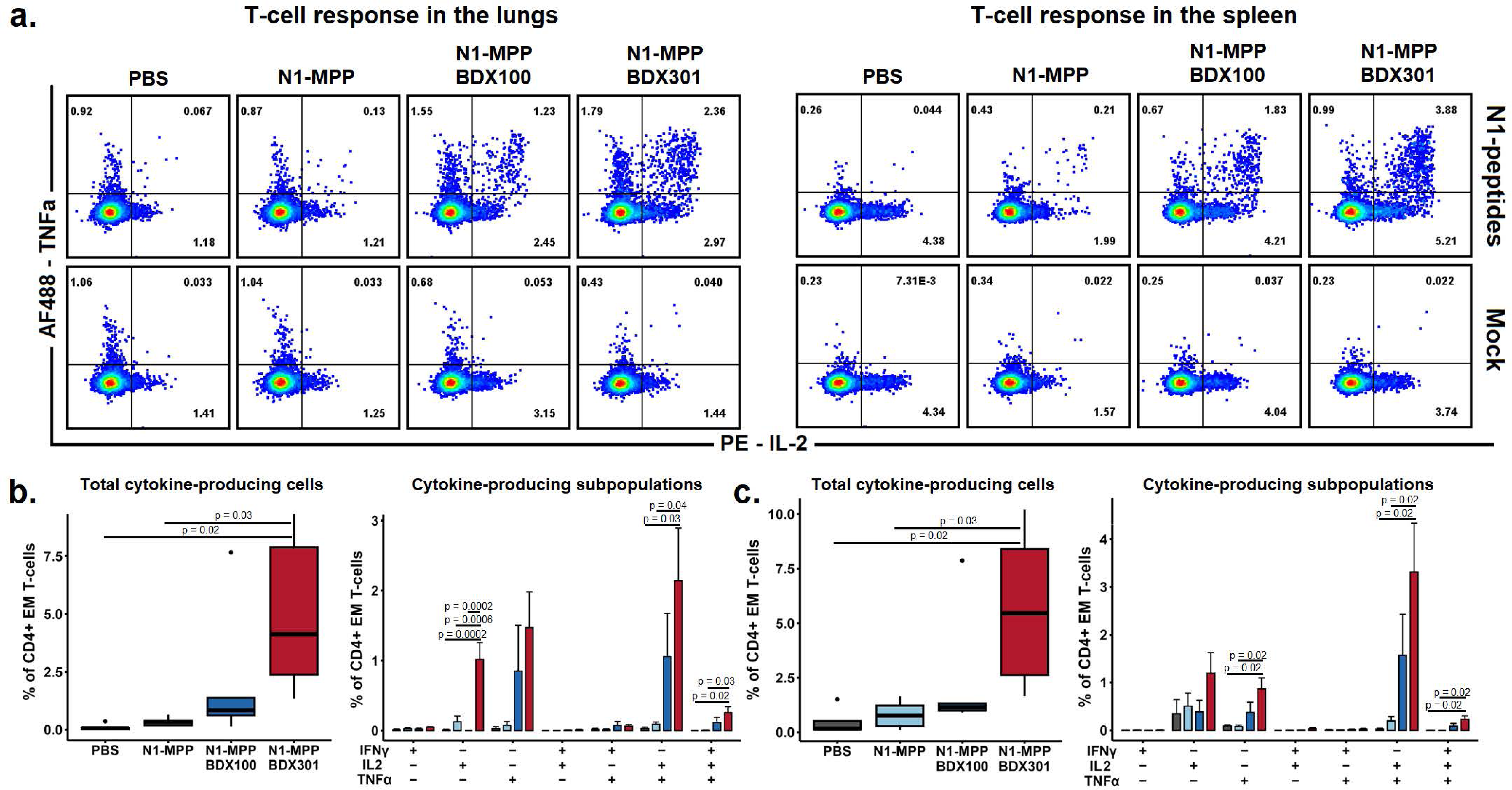
T-cellular immune response to the rNA-N1-MPP vaccine adjuvanted with BDX100 or BDX301. **a.** Representative flow cytometry plots showing the percentage of IL-2- and TNFα-producing CD4^+^ EM T-lymphocytes in mouse lungs and spleens 10 days after the booster immunization (gated on live CD3^+^CD19^‒^CD4^+^CD62L^‒^CD44^+^ cells). Cells were incubated with overlapping peptides, covering the whole sequence of N1 NA-protein for 6h in the presence of Brefeldin A and co-stimulatory anti-CD28 antibodies. **b.** Percentage of different cytokine-producing cell populations within the total CD4^+^ EM T-cell subset after background subtraction. Average and SE values are shown. Statistical analyses were performed using one-way ANOVA with Tukey posthoc test. P-values for the groups with statistically significant differences are shown on the plots.

### 3.4. BDX100 and BDX301 adjuvants induce partial cross-protection and stimulate T-cellular immune response to the heterologous challenge with A/bald eagle/Florida/W22-134-OP/2022 H5N1

To evaluate the cross-protective efficacy of the rNA-N1-MPP vaccine adjuvanted with BDX100 or BDX301, immunized BALB/c mice were challenged with 5 × LD_50_ of either the A/bald eagle/Florida/W22-134-OP/2022 H5N1 or prepandemic seasonal A/New Caledonia/20/1999 H1N1 influenza virus strains on Day 30 post-secondary immunization. Survival and body weight dynamics were monitored for 14 days post-challenge. For viral replication analysis, a subset of mice was challenged with 0.1 × LD_50_ of the same virus strains, and lungs and nasal turbinates were collected on Day 5 post-challenge to determine viral loads using a plaque assay.

Mice challenged with the A/bald eagle/Florida/W22-134-OP/2022 H5N1 virus experienced significant weight loss, but most animals in the adjuvanted vaccine groups survived, unlike those in the control and non-adjuvanted groups, where all mice succumbed to infection by Day 7. Recovery in the BDX100-adjuvanted group was marginally faster than in the BDX301 group. However, the rNA-N1-MPP vaccine did not confer cross-protection against the pre-pandemic seasonal A/New Caledonia/20/1999 H1N1 strain; most mice in both BDX100- and BDX301-adjuvanted groups succumbed to infection, as in the control group (Fig. 6a).

**Figure 6.**
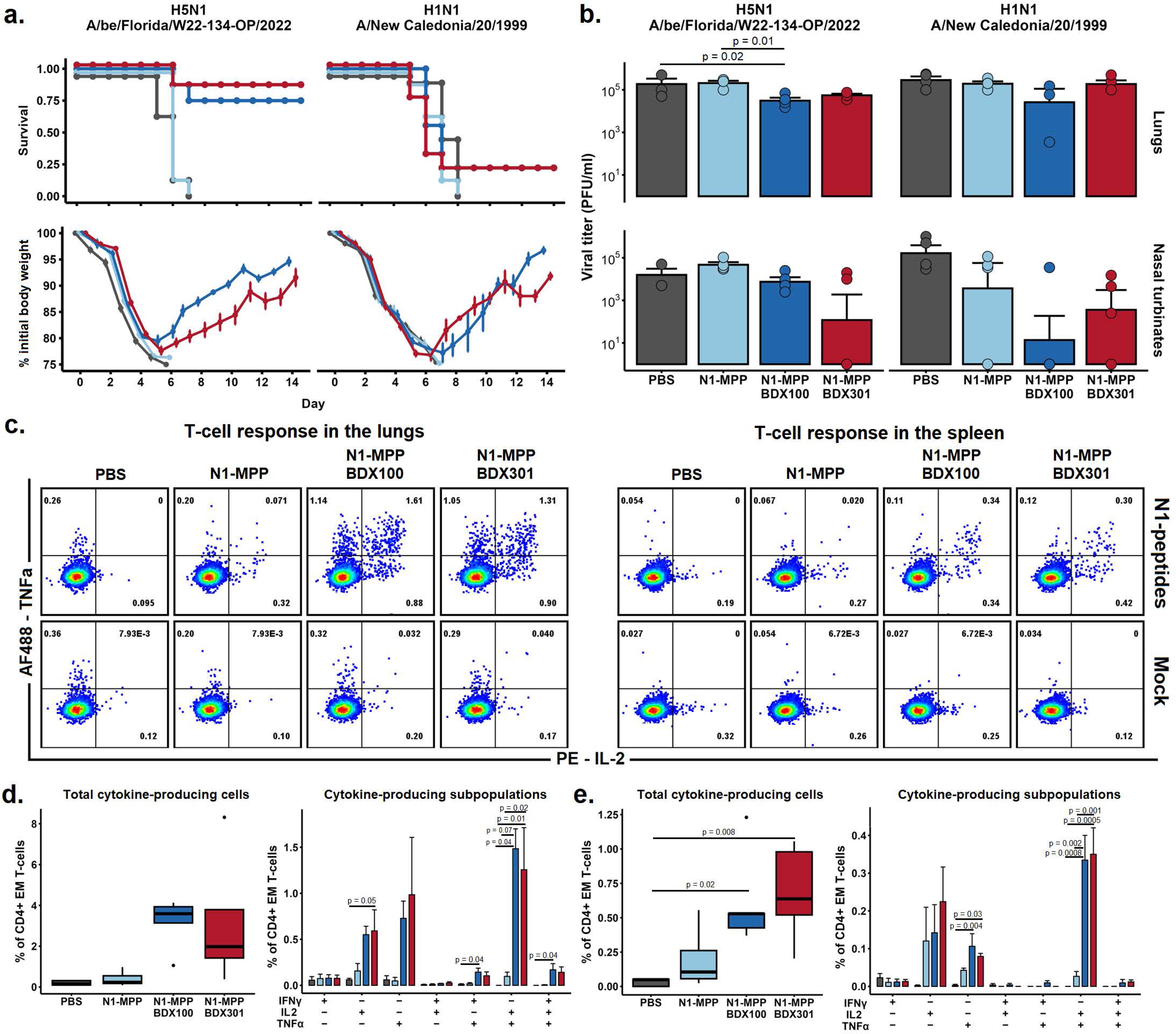
Protection and T-cellular immune response after the heterologous challenge. **a.** Body weight dynamics and survival after the heterologous challenge with 5 × LD_50_ of A/bald eagle/Florida/W22-134-OP/2022 (H5N1 reassortant with A/Puerto Rico/8/1934 vaccine backbone) and A/New Caledonia/20/1999 (H1N1) (n = 8). **b.** Viral titers in lungs and nasal turbinates on day 4 after the challenge (n = 4). **c.** Representative flow cytometry plots showing the percentage of IL-2- and TNFα-producing CD4^+^ EM T-lymphocytes (gated on live CD3^+^CD19^‒^ CD4^+^CD62L^‒^CD44^+^ cells) in mouse lungs and spleens 5 days after the challenge with 0.1 × LD_50_ of A/bald eagle/Florida/W22-134-OP/2022 (H5N1). Cells were incubated with overlapping peptides, covering the whole sequence of N1 NA-protein for 6h in the presence of Brefeldin A and co-stimulatory anti-CD28 antibodies. **d.** Percentage of different cytokine-producing cell populations within the total CD4^+^ EM T-cell subset after background subtraction. Average and SE values are shown. Statistical analyses were performed using one-way ANOVA with Tukey posthoc test. P-values for the groups with statistically significant differences are shown on the plots.

A significant reduction in viral replication was observed in the lungs of mice vaccinated with the BDX100-adjuvanted rNA-N1-MPP after challenge with the A/bald eagle/Florida/W22-134-OP/2022 H5N1 virus. However, vaccination did not affect viral replication in the upper respiratory tract or the replication of A/New Caledonia/20/1999 H1N1 in either the lungs. A tendency toward a decrease in the viral loads was shown in nasal turbinates of mice immunized with BDX100-adjuvanted rNA-N1-MPP after the challenge with A/New Caledonia/20/1999 H1N1 (Fig. 6b).

To assess the T-cell-mediated immune response following heterologous challenge, immunized C57BL/6 mice were challenged with 0.1 × LD_50_ of the A/bald eagle/Florida/W22-134-OP/2022 H5N1 virus. BALFs, lungs, and spleens were collected on Day 5 post-challenge, and extracted cells were restimulated with overlapping NA-derived peptides. Cytokine production was assessed using flow cytometry. Consistent with pre-challenge data, the antigen-specific response in vaccinated animals was dominated by effector-memory (EM) CD4^+^ T cells. Both BDX100 and BDX301-adjuvanted groups exhibited robust polyfunctional T-cell responses mediated by IFNγ^‒^ IL2^+^TNFα^+^ and IFNγ^+^IL2^+^TNFα^+^ subpopulations in the lungs and spleens. In contrast, the non-adjuvanted rNA-N1-MPP group showed only a marginal increase in antigen-specific single-positive T cells (IL2^+^ or TNFα^+^) without statistically significant differences compared with the control group (Fig. 6c, d).

To gain a deeper understanding of the protective mechanisms underlying post-vaccination immunity induced by the adjuvanted rNA-N1-MPP vaccine, we investigated the lung innate immune cell subpopulations, cytokine responses, and viral titers in bronchoalveolar lavage fluids (BALFs) of C57BL/6 mice infected with 0.1 × LD_50_ of the A/bald eagle/Florida/W22-134-OP/2022 H5N1 influenza virus on Day 5 after the challenge. Innate immune cell types, including resident and inflammatory monocytes, interstitial macrophages, alveolar macrophages, exudative macrophages, neutrophils, eosinophils, basophils, mast cells, dendritic cells, and NK cells, were identified using the protocol described by Yu *et al.*^69^. An additional group of unvaccinated and unchallenged animals (Intact) was included as a reference for baseline subpopulation values.

The infection triggered an influx of macrophages, monocytes, and neutrophils into the lung tissue in both vaccinated and non-vaccinated groups compared with intact mice. Notably, vaccinated groups receiving adjuvanted formulations exhibited reduced levels of inflammatory monocytes and NK cells relative to the rNA-N1-MPP and PBS groups, with the most substantial reductions observed in the BDX100-immunized mice. Conversely, neutrophil levels were significantly elevated in the lung tissue of adjuvanted vaccine recipients. The rNA-N1-MPP group showed a trend toward increased percentages of eosinophils, basophils, and mast cells compared with PBS or adjuvanted groups. All vaccinated groups demonstrated elevated levels of interstitial macrophages and dendritic cells relative to PBS control and intact mice (Fig. 7c). These cellular changes were accompanied by reduced viral titers in vaccinated groups, with the most pronounced reduction observed in the rNA-N1-MPP-BDX100 group, corroborating previously reported findings in BALB/c mice (Fig. 7d).

**Figure 7.**
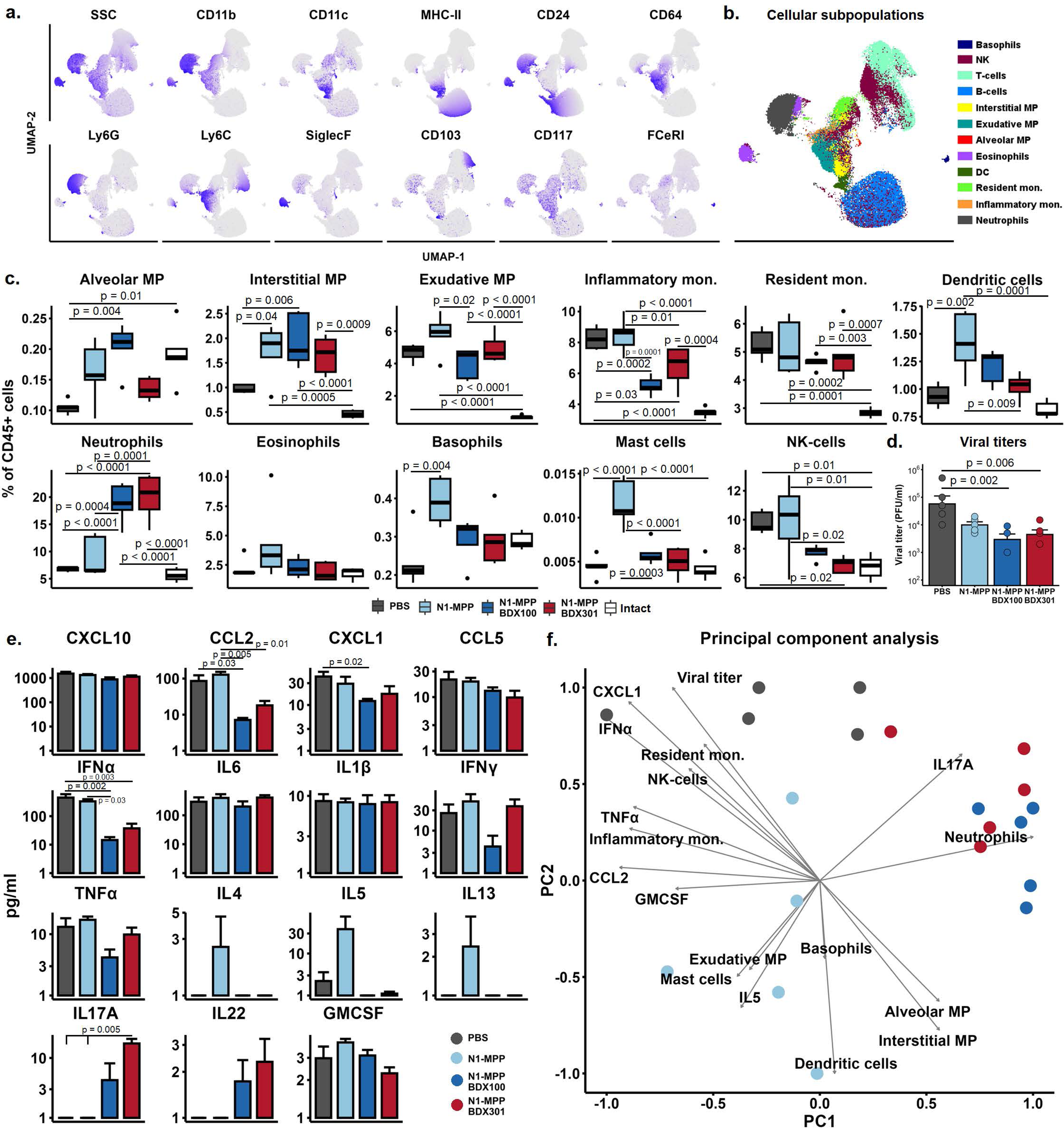
Innate immune response in mice vaccinated with rNA-N1-MPP vaccine adjuvanted with BDX100 or BDX301 after the challenge with A/bald eagle/Florida/W22-134-OP/2022 (H5N1) influenza virus. **a.** UMAP projections showing the differential expression of innate immune cells phenotype markers on lung cells. **b.** UMAP projection showing different innate immune cell subpopulations. **c.** Percentage of different innate immune cell populations in mouse lungs 5 days after the challenge with A/bald eagle/Florida/W22-134-OP/2022 (H5N1 reassortant with A/Puerto Rico/8/1934 vaccine backbone) influenza virus (mean ± SE). **e.** Cytokine concentrations in BALFs on day 5 after the challenge (mean ± SE). Statistical analyses were performed using one-way ANOVA with Tukey posthoc test. P-values for the groups with statistically significant differences are shown on the plots. **f.** Principal component analysis (PCA) biplot represents the distribution of mice in accordance with the parameters of the immune response as well as factor loads of analyzed variables on the first two principal components. Variables for the PCA were selected based on the results of group comparison using ANOVA.

Vaccination with adjuvants also decreased the levels of pro-inflammatory cytokines and chemokines, including CCL2, CXCL1, IFNα, IFNγ, and TNFα, in the BALFs of infected mice. However, the BDX-adjuvanted groups exhibited increased production of IL17A and IL22, consistent with elevated neutrophil levels. In contrast, the rNA-N1-MPP group showed enhanced concentrations of Th2-associated cytokines IL4, IL5, and IL13 in response to the challenge (Fig. 7e).

Principal component analysis (PCA) provided an integrated view of the immune response on Day 5 post-challenge with 0.1 × LD_50_ of H5N1. Statistically significant variables identified by ANOVA were used to generate the PCA biplot (Fig. 7f). The analysis revealed that BDX100 and BDX301-adjuvanted groups exhibited similar immune response profiles to the heterologous challenge. These profiles were distinctly different from those of the rNA-N1-MPP and PBS control groups, primarily driven by elevated IL17A levels and neutrophil percentages in the lungs. Conversely, the non-adjuvanted vaccine elicited partial Th2 polarization, characterized by increased basophils and mast cells in the lungs and elevated IL5 production.

## 4. Discussion

Recombinant NA-proteins are actively studied as potential broadly protective influenza vaccines. It has been shown that intramuscular immunization with NA-based constructs provides efficient protection against homologous and heterologous influenza viruses in a diverse array of animal models including mice, hamsters, guinea pigs, rabbits^14^, pigs^70^, and ferrets.^27,43,53,56,71^. The potential use of NA as a mucosal vaccine, however, is under-investigated. Mucosal surfaces of the respiratory tract are in direct contact with the environment and serve as the entry point for many pathogens including the influenza virus. Activation of the immunological protective mechanisms of the mucosal surfaces through vaccination allows the prevention of the virus entry at the early stage of infection. It also limits pathogen shedding, restricting transmission in the population. It was shown previously in animal models^29,72^ and clinical studies^73–75^ that intranasal vaccination offers superior protection against the influenza virus compared with intramuscular immunization.

In the present study, BDX100 and BDX301 adjuvants significantly enhanced rNA-N1-MPP-induced protection compared with the non-adjuvanted vaccine. Animals vaccinated with the adjuvanted formulations maintained stable body weight and survival after the challenge with the homologous A/Singapore/GP1908/2015 H1N1 strain. Partial protection was observed against the heterologous A/bald eagle/Florida/W22-134-OP/2022 H5N1 strain, with reduced mortality and lung viral titers. The BDX100-adjuvanted vaccine led to slightly better outcomes than BDX301. However, neither adjuvant provided protection against the heterologous pre-pandemic seasonal A/New Caledonia/20/1999 H1N1 strain which is antigenically very distinct. Both adjuvants enhanced humoral immunity, increasing the level of anti-N1 cross-reactive IgG antibodies against recent H1N1 and H5N1 strains including a bovine H5N1 NA. Mucosal immune responses were also boosted, with increased IgA production and activation of GCBs and Tfhs, particularly in the upper respiratory tract. It was shown previously that intranasal, but not systemic immunization induces local IgA production in the lungs and the level of these antibodies correlates with protection against a challenge with homologous and heterologous viruses, providing superior immunological defense compared with the circulating antibodies alone^29^. IgAs in the lungs are secreted by local B-cell subsets that participate in the protection against a secondary challenge^76^.

The distinctive feature of the present study was the intranasal administration of the vaccine in a volume of 10 µl (5 µl/nare) which restricts the immunization predominantly to the upper respiratory tract with a low direct impact on lungs^77^. However, the adjuvanted formulations successfully triggered the mucosal immune response not only in URT but also in the lungs, inducing antibody production and altering the cellular subpopulation content. This finding reflects the integrative nature of mucosal immunity, which allows the generation of the immune response in the mucosal tissues anatomically distant from the site of antigen administration^78,79^.

In the URT, the two most affected cell populations were GCB cells and Tfh lymphocytes. Previous research using various influenza vaccine platforms in mice has demonstrated the role of germinal centers (GCs) in enhancing immune protection against influenza. GCB cells undergo proliferation, antibody class-switching, and affinity maturation in the GCs. This increases the antibody production and neutralizing responses^80^. Greater frequencies of GCB cells correlate with improved protection against influenza virus challenges^81–83^. The maturation of GCBs is controlled by the follicular T-helper lymphocytes which represent a specialized type of CD4^+^ T cell that reside in GCs within secondary lymphoid organs. Tfh cells provide essential signals to B cells within GCs through the co-stimulatory molecule CD40L and cytokines such as IL21, driving their differentiation into antibody-secreting plasma cells and long-lived memory B cells^84,85^. Tfhs are indispensable for the generation of an efficient immune response to influenza vaccination^86,87^. BDX100 and BDX301 adjuvants used in this study consist of TLR4 and TLR2 agonists. Both TLR4 and TLR2 can be expressed on Tfh cells and when stimulated increase the differentiation of Tfh cells and promote the secretion of IL21^88^. TLR4 and TLR2 are also expressed on DCs. The activation of these receptors can lead to increased expression of co-stimulatory molecules (such as CD40, CD80, and CD86) and the production of pro-inflammatory cytokines. These signals can promote the differentiation of naive CD4^+^ T cells into Tfh cells^89^.

rNA-N1-MPP adjuvanted with BDX100 or BDX301 induced strong systemic and local T-cellular immune response, mediated by the polyfunctional CD4^+^ T cells and associated with the accumulation of CD4^+^ and CD8^+^ T_RM_ cells in the lung tissue. It was shown previously that CD8^+^ T_RMs_ are required for optimal cross-protection against pulmonary virus infection^38^. In this study, despite the substantial increase in the total number of CD8^+^ T_RM_s in the lungs, the antigen-specific T-cellular immune response was shown only for CD4^+^ T-lymphocytes. The role of CD4^+^ T-cells in the mediation of the protective response to influenza virus is controversial. Some studies showed that CD4^+^ T cells play an indirect role in protection, stimulating an early antibody response, and cytokine production, providing signals for the recruitment of CD8^+^ T-lymphocytes and innate immune cells, and stimulating the differentiation of long-lived memory CD8^+^ T cells^90,91^. Memory CD4^+^ T cells can work in synergy with B cells stimulating the secretion of virus-neutralizing antibodies. In the absence of other lymphocytes, memory CD4^+^ T cells mediate antigen-specific viral clearance via IFNγ and perforin^92^. It was shown that the depletion of CD4^+^ T cells at the time of vaccination diminishes CD8^+^ T cell recall response. Adoptive transfer of CD4^+^ T cells did not lead to viral load decrease in contrast to CD8^+^ T-cell transfer but similarly resulted in increased survival^93^. In another study, mice vaccinated with FluMist demonstrated a 50% survival rate after CD4^+^ T-cell depletion with a subsequent lethal challenge, whereas after CD8^+^ T-cell depletion, the survival rate was only 25%^94^. In another study, however, the depletion of CD4^+^ T cells did not influence the protection after the challenge in whole inactivated virus (WIV)-vaccinated mice, even though IFNγ-producing CD4^+^ T cells were present after the vaccination^95^. Further studies are required to distinguish between the contribution of local antibody response, systemic humoral response, and T-cellular immune response to the overall protection against the homologous and heterologous challenges in rNA-N1-MPP vaccinated mice.

BDX100 and BDX301 adjuvants modulated the innate immune response to the heterologous virus infection by A/bald eagle/Florida/W22-134-OP/2022 H5N1 in rNA-N1-MPP-vaccinated mice. The most noticeable effect was associated with an increase in the concentration of IL17A accompanied by the accumulation of neutrophils in the lung tissue. IL17A is a cytokine primarily produced by Th17 cells, γδ T cells, and innate lymphoid cells. It plays a pivotal role in recruiting neutrophils to sites of infection by inducing the production of chemokines such as CXCL1 and CXCL2^96,97^. Elevated levels of IL17A after heterologous challenge suggest that the vaccine-induced immune response favors a Th17-skewed pathway. Neutrophils are frontline innate immune responders, critical for early control of viral infections. In the context of influenza, neutrophils can help clear virus-infected cells and provide an early antiviral response, but excessive neutrophil activity might also contribute to immunopathology and tissue damage^98^. It was shown previously that mucosal routes of vaccination favor the generation of Th17 cells^99–101^ which can lead to excessive inflammation and higher morbidity upon a secondary influenza virus infection^102^. In this study, however, the production of IL17A and the accumulation of neutrophils in the lungs was not associated with excessive immunopathology since mice immunized with BDX100/BDX301-adjuvanted vaccine demonstrated higher survival and lower morbidity after homologous and heterologous challenge. Influenza virus is known to suppress neutrophil functions, which can lead to bacterial superinfection – a common complication of influenza virus infection^103,104^. Increased IL17 production correlates with enhanced neutrophil infiltration and improved survival against bacterial superinfection^105^. Thus, the findings presented in this study suggest that BDX100 and BDX301 adjuvants can potentially be beneficial for the prevention of bacterial superinfection during the postvaccinal immune response to an antigenically mismatched strain. This hypothesis will be evaluated in future research.

The present study highlights the promising potential of N1-MPP influenza vaccines adjuvanted with BDX100 or BDX301 for inducing robust mucosal and systemic immunity. Both adjuvants significantly enhanced vaccine efficacy, with BDX100 showing marginally superior results. These results pave the way for further research to optimize adjuvanted NA-based vaccine formulations for clinical use.

### Limitations of the study

As this study was performed in inbred naïve mice, it is difficult to account for factors such as pre-existing immunity to influenza viruses/antigenic imprinting, when considering a clinical application of NA vaccines in humans. Antigenic imprinting has different effects on NA versus HA immunity, but could result in dampened anti-NA immune responses induced by vaccination if the NA is co-administered with an HA previously encountered by the immune system^106,107^. Experiments with pre-immune animals may be useful for optimizing NA antigen choice in a vaccine intended for humans. As the mice in our study had no pre-existing immunity and NA was the only viral component of the vaccine, only NA-specific T-cell responses were evaluated. In humans, however, most T-cell responses associated with protection are directed against more conserved internal proteins, particularly the nucleoprotein, whereas the contribution of NA-specific T-cell responses to protection is not well known. NA-based vaccines may however promote back-boosting of responses to more conserved viral proteins following exposure to drifted viral strains or non-targeted subtypes. This was demonstrated by Choi et al., for example, who vaccinated pre-challenged mice and found that infection-permissive vaccination strategies better boosted memory T cells and promoted cross-protective CD8^+^ cellular-mediated responses to IAV, than HA-targeting strategies^108^. Experiments using mice with pre-existing influenza immunity, established by viral challenge or different vaccinations, may be useful for addressing these questions.

Mice are also an imperfect model for intranasal vaccination evaluation, particularly as variations in the administered volume of vaccine, and differences in their mucosal environment may have a more significant effect on the types of immune responses being evaluated. Additionally, the potential for an intranasal vaccine to inhibit viral transmission cannot be evaluated in mice, as they do not efficiently transmit influenza virus in laboratory conditions. Differences in types of TLRs and their expression between mice and humans should also always be considered for adjuvant evaluation. Further experiments in alternative animal models with larger respiratory tracts and immunology/TLR expression closer to those of humans, such as the ferret model, could address these limitations. Experiments using less expensive small animal models for IAV, such as guinea pigs or hamsters, would also be valuable to evaluate transmission inhibition and would also help mitigate the substantial effect variation in administration volume has on intranasal vaccination observed in mice.

## Acknowledgements

This study was supported by the Collaborative Influenza Vaccine Innovation Centers (CIVIC) contract 75N93019C00051 (F.K.).

---

**Supplementary figure 1.**
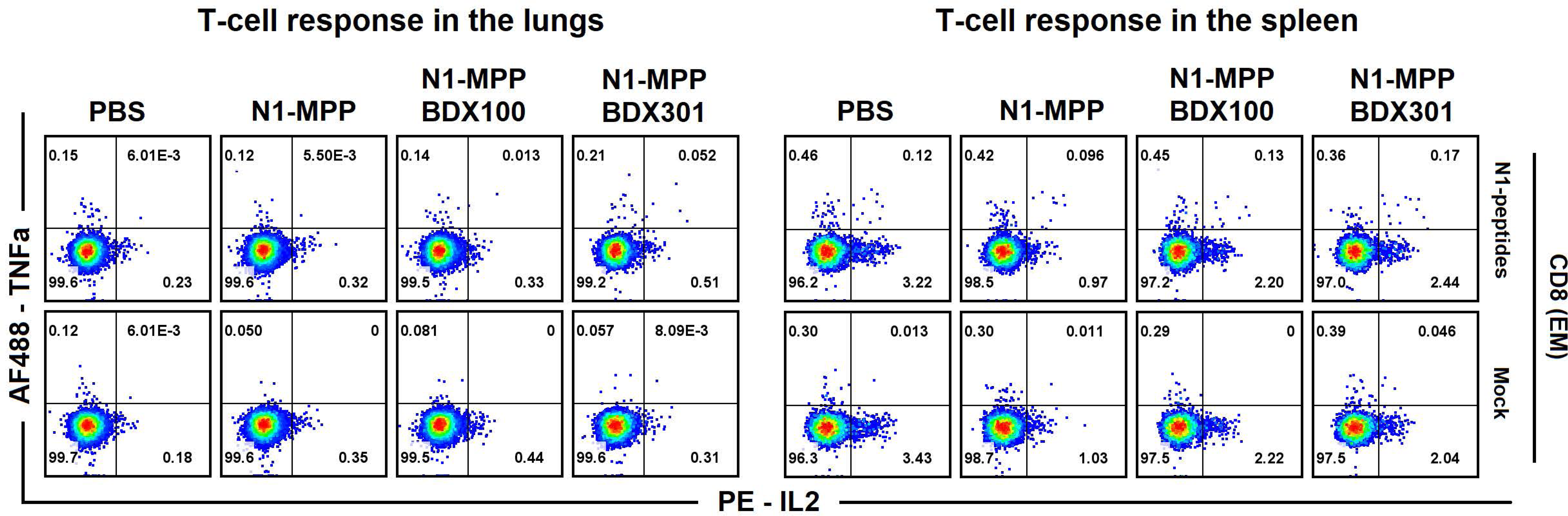
CD8^+^ T-cellular immune response to the N1-MPP vaccine adjuvanted with BDX100 or BDX301. Flow cytometry plots showing the combined data on the percentage of IL-2- and TNFα-producing CD8^+^ EM T-lymphocytes in mouse lungs and spleens 10 days after the booster immunization. Cells were incubated with overlapping peptides, covering the whole sequence of N1 NA-protein for 6h in the presence of Brefeldin A and co-stimulatory hanti-CD28 antibodies.

